# Small-angle x-ray microdiffraction from fibrils embedded in tissue thin sections

**DOI:** 10.1101/2022.05.10.491381

**Authors:** Prakash Nepal, Abdullah Al Bashit, Lin Yang, Lee Makowski

## Abstract

Small-angle x-ray scattering (SAXS) from fibrils embedded in a fixed, thin section of tissue includes contributions from the fibrils; the polymeric matrix surrounding the fibrils; other constituents of the tissue; and cross-terms due to the spatial correlation between fibrils and neighbouring molecules. This complex mixture severely limits the amount of information that can be extracted from scattering studies. However, availability of micro- and nano-beams has made possible measurement of scattering from very small volumes which, in some cases, may be dominated by a single fibrillar constituent. In those cases, information about the predominant species may be accessible. Nevertheless, even in these cases, the correlations between the positions of fibrils and other constituents have significant impact on the observed scattering. Here, we propose strategies to extract partial information about fibril structure and tissue organization on the basis of SAXS from samples of this type. We show that the spatial correlation function of the fibril in the direction perpendicular to the fibril axis can be computed and contains information about the predominant fibril structure and the organization of the surrounding tissue matrix. It has significant advantages over approaches based on techniques developed for x-ray solution scattering. We present examples of the calculation of correlation in different types of samples to demonstrate the kinds of information that can be obtained from these measurements.

**Synopsis:** The availability of micro- and nano- x-ray beams is making possible measurement of scattering from very small volumes, opening possibilities for deriving *in situ* structural information on fibrillar constituents in complex materials and tissues. This work outlines a set of strategies for confronting the formidable technical obstacles to extracting useful structural information from scattering derived from these materials.

## 1. Introduction

Understanding the molecular basis of developmental or disease processes may require detailed information about the distribution of constituents throughout a tissue. For instance, immuno-histochemistry can be used to map the distribution of specific epitopes within tissues. However, methods based on optical microscopy often have inadequate resolution for the questions being asked and techniques that more deeply probe the molecular structure of constituents are required. X-ray scattering from thin sections of tissue is rarely attempted because the heterogeneous mixture of constituents makes data interpretation difficult, the disruption of native structure intrinsic to sample preparation may destroy relevant structural features and the disorientation and disorder of structures that do survive weaken and obscure the collected data. However, availability of micro- and nano-beams make possible measurement of scattering from very small volumes which, in some cases, may be populated largely by a single constituent. Furthermore, some classes of relevant structures, including pathological fibrillar protein deposits implicated in Alzheimer’s and other neurodegenerative diseases have robust architectures that are particularly resilient to the physical and chemical insults of sample preparation. The impact of formaldehyde fixation and ethanol dehydration is to create a rigid, cross-linked polymer matrix. There have been extensive studies on the chemistry of formaldehyde reactions with amino acids and proteins (French and Edsall, 1945; Werner et al., 2000), the impact on availability of epitopes for immunostaining (Werner et al., 2000) and its effect on FTIR spectra (Zohdi et al., 2015). These studies have shown that fixation preserves most protein secondary structures (Zohdi et al., 2015) and epitopes (Werner et al., 2000). Neuropathological fibrils stabilized by a core of cross-β structure appear to be highly resilient to these treatments and are left intact, trapped within the cross-linked matrix (Liu et al., 2016). Of particular importance to analysis of SAXS data, ethanol dehydration alters the electron density contrast between scattering particles and their surroundings and removes essentially all lipids (Zohdi et al., 2015).

In many cases the fibrillar structures of interest are ‘frozen’ with random orientation in the matrix of fixed tissue and scattering from these fibrils might be considered analogous to solution scattering of fibrils in aqueous solution. However, the presence of a cross-linked macromolecular network in which the fibres are embedded presents significant challenges for analysis of scattering data. In this paper we present an analysis of the properties of scattering from fibrillar objects embedded within a polymeric matrix and provide strategies for extracting information from these data. The methods described are of particular relevance to analysis of pathological protein deposits in human brain tissue.

SAXS is routinely used to obtain the shape and size of globular molecules in solution (Glatter and Kratky, 1982; Svergun and Stuhrmann, 1991; Svergun and Koch, 2003; Koch, Vachette and Svergun, 2003; Pollack, 2011; Putnam et.al., 2007; Lattman et al., 2018). But conventional methods for analysis of SAXS data are not well suited to deal with fibrils that have very long axial ratios; or scattering particles embedded in a polymeric matrix. SAXS data are frequently used to calculate the pair distribution function (Svergun, 1992; Liu and Zwart, 2012; Hong and Hao, 2009) - a histogram of the interatomic vector lengths in a scattering particle. However, the pair-distribution function is less informative when the scattering particle is a fibril of indeterminant length: The maximum interatomic vector length routinely obtained from the pair-distribution function is no longer well defined. In these cases, use of cross-section scattering functions is preferred (Porod, 1948; Kratky and Porod, 1948; Guinier and Fournet, 1955; Kratky, 1963; Glatter, 1980). Here we demonstrate the utility of this approach, possible when equatorial scattering can be isolated from off-equatorial scattering. In particular, when the axial repeat distance of a fibril is less than ∼ 30 Å, small angle intensity may be due entirely to equatorial scattering (as defined by conventional fiber diffraction methods) and the equatorial intensity can be derived from the observed intensity by a simple geometric correction (Makowski, 1978). The resulting equatorial intensity can be used to calculate a ‘cross-section pair-distribution function’ (Glatter, 1980), analogous to the conventional pair-distribution function, but representing a histogram of the interatomic distances projected onto the equatorial plane of the fibril. The length of the scattering fibril is irrelevant for this calculation.

A second change from conventional SAXS analysis is a focus on the correlation function (rather than the pair-distribution function). This has the advantage that the correlation function is much less sensitive to errors in measurement of intensity at very small scattering angles or the absence of intensity measurements very near zero-angle. The correlation function is the probability that, given there is an atom at some origin position there is also one at a distance r. As is the case for the pair-distribution function, the correlation function can be defined for conventional (spherical) SAXS analysis or for the fibrous (cross-sectional) case where the relevant distances are those perpendicular to the fiber axis.

Here, we start with an overview of sample preparation and fiber diffraction theory and then present examples of scattering from fibrils in environments of increasing complexity. Starting with calculation of scattering from solid cylinders, we progress to scattering from molecular models of fibrils; analysis of scattering from fibrils in aqueous solution; in concentrated gels and finally to fibrils embedded in tissue. This progression allows us to demonstrate the impact of the milieu in which the structures are immersed on the properties observed.

## 2. Methods

### 2.1. Sample preparation

Data from tissue samples described here were collected from samples prepared at the Massachusetts Alzheimer’s Disease Research Center (MADRC) at the Massachusetts General Hospital using standard neuropathological processes to produce 20 μ sections with chemical and physical properties typical of histological sections as previously described (Liu et al., 2016). These unstained sections are thicker than conventional histological sections in order to increase the volume of material irradiated. A 5 μ beam was scanned over a rectangular region of interest (ROI) chosen on the basis of examination of serial sections stained for Aβ and tau proteins. Regions exhibiting high levels of Aβ or tau were chosen for examination. Tissue sections were spread on either 1 cm x 1 cm mica films (12 μ thick) or 2.7 × 2.7 mm^2^ SiN membranes (1 μ thick) and then mounted on sample holders printed to LiX specifications (see below). These holders were mounted directly on the LiX beam line stage (Yang et al., 2020; 2022).

### 2.2. Data Collection

Tissue sections mounted on mica or SiN films were scanned with a 5 μ x-ray microbeam using 5 μ steps to collect diffraction patterns as a function of position on a square grid across the tissue section. Data collection was carried out at the LiX beam line at the NSLS-II synchrotron source at Brookhaven National Laboratory. Scans were carried out over a rectangular area of between 300 × 300 μ ^2^ (3600 diffraction patterns) and 600 × 600 μ ^2^ (14,400 diffraction patterns). An exposure time of 0.5 seconds was used and (including data transfer and sample step) approximately 0.8 seconds was required per exposure, or about 48 minutes for a 300×300 μ ^2^ region of interest (ROI). Data were collected on SAXS and WAXS (wide-angle x-ray scattering) detectors simultaneously (see Figure 1), circularly averaged and merged using LiX-specific software.

**Figure 1.**
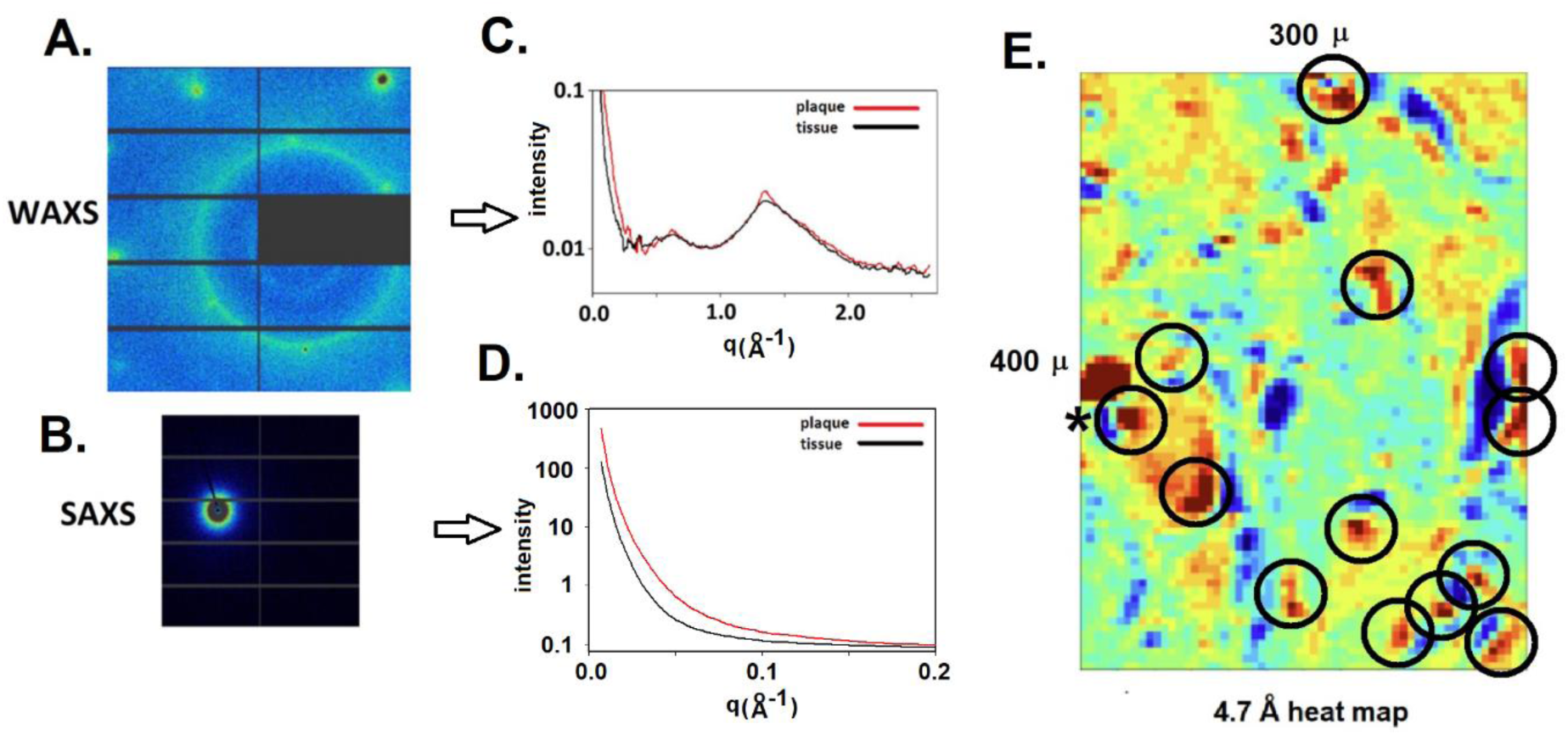
X-ray data from a histological thin section of brain tissue. Scattering data are collected simultaneously on SAXS and WAXS detectors (A). Scattering from mica substrate is subtracted and the resulting intensities circularly averaged and merged (B). Tissue scattering is subtracted from lesion scattering, resulting in estimate of scattering from constituent fibrils (C). The scattering at 4.7 Å spacing is prototypical of cross-β structures including those made from Aβ or tau. Maps of intensity at 4.7 Å spacing as measured in 4941 scattering patterns in a region of interest (D) make it possible to locate lesions (circles).

Data from *in vitro* assembled Aβ fibrils and TMV particles in solution used in the calculations described here have been reported previously (Roig Solvas and Makowski, 2017). Data from concentrated gels of TMV particles are from Caspar (1955).

### 2.3. Data Processing

Intensities in diffraction patterns were pre-processed to remove scattering from the mica films used as substrates for the tissue. This involved the ‘masking’ of peaks due to mica using a modification of LiX-specific software (Yang et al., 2020). Three of these peaks can be seen near the left and top edges of the diffraction pattern in **Figure 1A**. After masking, data from the SAXS and WAXS detectors (**Figures 1A and 1B**) were circularly averaged, merged and scaled using LiX-specific software, resulting in intensity distributions such as those in **Figure 1C and 1D**. All intensities were corrected for small fluctuations in beam intensity as measured at the LiX beam line prior to further processing. The location of pathological protein deposits could usually be determined on the basis of intensity observed at q ∼ 1.34 Å^-1^ (corresponding to a spacing of 4.7 Å). This intensity was calculated from circularly averaged data for all patterns in an ROI and displayed as a ‘heat map’ such as the one in **Figure 1E**, showing the distribution of relative intensities at a spacing of 4.7 Å in data from 4941 (61×81) scattering patterns in a 300 × 400 μ^2^ ROI. Subtraction of tissue scatter from scattering observed in lesions results in an estimate of scattering from fibrils (**Figure 1F**) because cross-terms due to correlation of fibril positions with surrounding tissue constituents are generally insignificant in the WAXS regime.

Correlation functions were calculated using an indirect Fourier transform (Glatter, 1980) with some modifications. The indirect Fourier transform as utilized by Glatter (1980) and Svergun (1992) requires knowledge of the maximum length of interatomic vectors in a scattering particle. In scattering from very long fibrils, that maximum length provides little constraint on the calculation. Furthermore, when embedded in a tissue matrix, very long interatomic correlations may be present. In many cases those correlations may give rise to scattering at sufficiently small angles as to preclude their measurement. In such cases, the maximum correlation distance observable is dictated by q_min_. The maximum interatomic vector length used in the indirect transform is then chosen to be (2π/q_min_) rather than d_max_. This results in calculation of correlation functions that represent a lower bound on correlations within and among particles in the scattering volume.

### 2.4. Background subtraction

Lesions commonly found in brain tissue in Alzheimer’s disease contain high concentrations of fibrillar structures made up of either Aβ peptides that are the primary constituent of ‘plaques’ or tau proteins that constitute the predominant constituent of ‘neurofibrillary tangles’. Other tissue constituents permeate the lesions, surround the fibrils and contribute to the observed scattering. Usually, the overall intensity distribution in scattering from lesions is very similar to that observed from tissue adjacent to the lesions. But it is distinguished by the presence of a sharp peak at 4.7 Å spacing arising from the cross-β spine of the fibrils. Although the diffuse scattering from a lesion is similar in overall shape to that from adjacent tissue, it is usually somewhat more intense. This increased intensity appears to be due to greater density of material within the lesion compared to adjacent tissue (Nepal et al., 2018). Scattering from the non-fibrillar tissue constituents is removed from fibrillar scattering within the plaque by assuming that the scattering of the tissue constituents within the lesion is the same as in a neighboring region devoid of fibrils. Since scattering from the underlying mica substrate does not require scaling, we carried out subtraction according to I = (I_l_ - I_b_) - a (I_t_ - I_b_) where I_l_ is the observed scattering from the lesion; I_t_ that observed from proximal tissue; I_b_ background observed from scattering from mica devoid of tissue (Bashit et al., 2022). The scale factor, a, is chosen to minimize the difference between plaque and background intensities in the range 1.6 < q < 2.0 Å^-1^, where all evidence has indicated the fibrillar scattering is weak.

### 2.5. Scattering from tissue

A thin section of fixed human brain tissue contains a network of cross-linked macromolecules, largely dehydrated and adhered to a solid substrate. In tissue from Alzheimer’s subjects these sections contain occasional lesions (plaques or tangles) in which pathological fibrillar structures may be the predominant constituent (**Figure 2**). When the constituents exhibit essentially no orientation, the (circularly symmetric) scattering from these samples is given by the Debye formula,

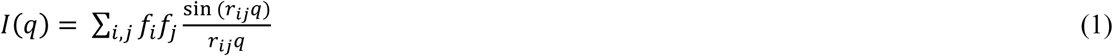

where the sum is over all atoms in the scattering volume, r_ij_ is the distance between atom i and atom j, *f*_i_ is the scattering factor of atom i and q is the momentum transfer (q = 4πsin(θ)/λ where 2θ is the scattering angle and λ is the wavelength of the incident x-rays). In scattering from the lesions embedded within the tissue sample, there will be contributions from fibrils, other tissue constituents and the support substrate (mica film):

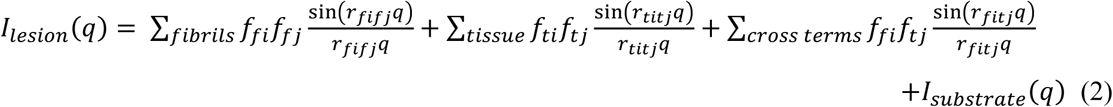

where f_fi_ represents the scattering factor of an atom within the fibrils, f_ti_ of atoms within the tissue and *I*_*substrate*_(q) is the background from mica substrate. The physical separation between substrate and lesion or tissue means that cross-terms with substrate correspond to interatomic vectors longer than ∼ 1000 Å and will not measurably contribute to observed scattering. If the structure of the tissue network within the lesion is essentially identical to that outside the lesion, the second and fourth terms on the right side of equation (2) can be removed by subtracting intensity observed from a lesion-free region of the tissue. However, the cross-terms between fibrils and tissue are not removed by this subtraction. If the molecular environment within the tissue were completely unstructured, as is the case outside the hydration shell of macromolecules in aqueous solution, the cross-terms would contribute relatively little to the scattering (Park et al., 2009) and could be, to a large extent, ignored. However, this is not the case for fixed tissue. Therefore, we can calculate directly from observations:

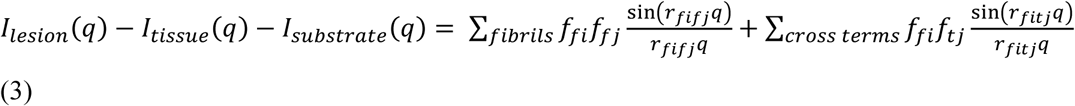

**Figure 2.**
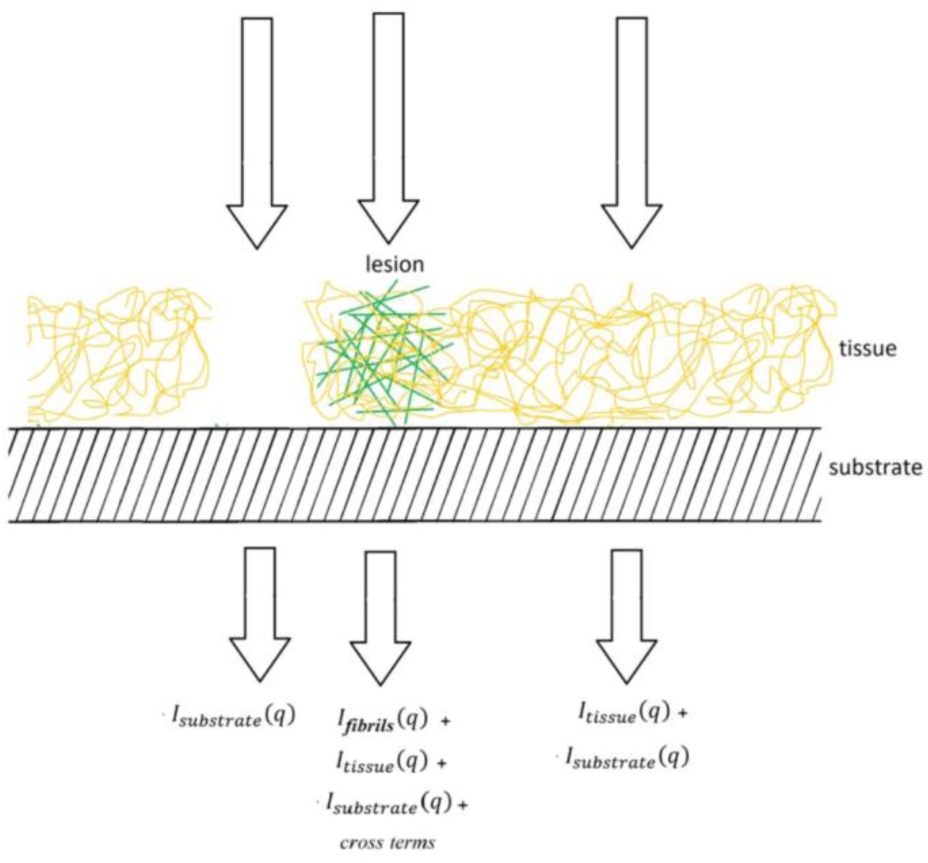
Diagram of tissue sample mounted on solid substrate and the types of data that can be collected.

The first term on the right-hand side of equation (3) contains information about the structure of individual fibrils and the second term information about correlation in the positions of tissue constituents with the fibrils. A partial separation of the contribution of these two terms may be possible since the first term is dominated by short interatomic vectors; the second term by longer vectors.

### 2.6. Fiber diffraction

Fibrous biological assemblies are usually helical arrangements of identical subunits, the Fourier transform of which is a series of layer planes perpendicular to the fiber axis, arranged at spacings reciprocal to the axial repeat of the fiber (**Figure 3**). Cross-β structures such as amyloid fibrils are composed of stacked Aβ peptides with an axial spacing of ∼ 4.7 Å, and the intensity in corresponding diffraction patterns is limited to layer planes spaced at 1/4.7 Å^-1^, the first order of which is frequently observed in diffraction patterns from these structures (**Figure 1**). In tissue these fibrils exhibit little or no preferred orientation and give rise to scattering patterns that are circularly symmetric, or nearly so, as diagrammed in **Figure 3**. These patterns can be divided into the intensity at spacings smaller than 1/4.7 Å^-1^, which is due to intensity in the equatorial plane of the fiber pattern; and intensity at q ≥ 1/4.7 Å^-1^ that will be dominated by scattering from the first layer plane. Analysis of the equatorial scattering can provide information about the diameter and shape of the fibrils and the arrangement of material around the fibril within the tissue (Roig Solvas and Makowski, 2017). Comparison of the shape of the 4.7 Å reflection with that predicted from high resolution structures obtained by cryoEM studies of isolated or *in vitro*-assembled material may provide identification of the precise structure within the scattering volume (Bashit et al., 2022). In this report, we limit our analysis to the small-angle regime.

**Figure 3.**
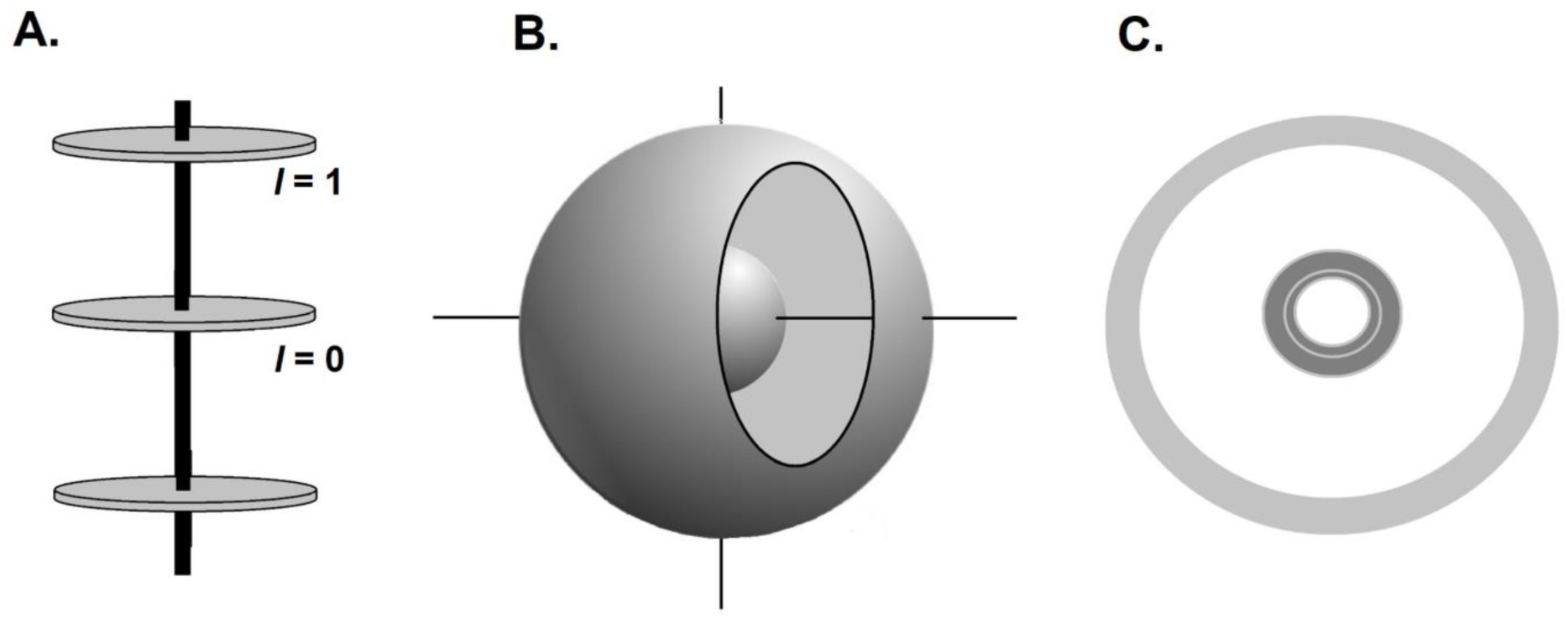
(A) Diffraction from a periodic, fibrous structure is limited to layer planes spaced at distances reciprocal to the axial repeat of the fiber. Equator is indexed as *l* = 0; first layer line, *l* = 1 and so on. For a cross-β structure the axial repeat is ∼ 4.7 Å. When the fibrous structures are completely disoriented, the fiber pattern is spherically averaged in reciprocal space (B), giving rise to a diffraction pattern that is circularly symmetric (C). The disorientation spreads the two-dimensional layer planes (including the equator) onto three-dimensional surfaces.

## 3. RESULTS

### 3.1. Calculation of scattering from disoriented solid cylinders

Interpretation of the small-angle scattering from disoriented fibrils requires consideration of the impact of disorientation of the fibrils; the finite and often heterogeneous and indeterminant length of the fibrils; and the impact of components of the surrounding tissue on scattering from the fibrils. At sufficiently low resolution, fibrils roughly approximate solid cylindrical structures. The Fourier transform of a solid cylinder of radius r_cyl_ and length h is (Oster and Reilly, 1952):

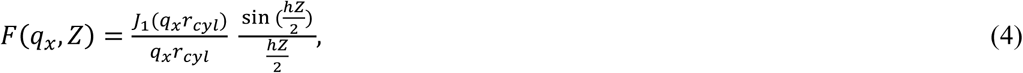

where q_x_ is the distance from the cylindrical axis in reciprocal space. For infinitely long cylinders the three-dimensional Fourier transform is confined to the equator (Z = 0) and equation (4) reduces to that for Fraunhofer diffraction by a circular aperture first derived by Airy (Airy,1835). Scattering from randomly oriented solid cylinders exhibiting no correlation of spatial positions can be readily computed from equation (4) and that intensity can be used to calculate the pair-distribution function, P_s_(r) (Svergun and Koch (2003)) according to

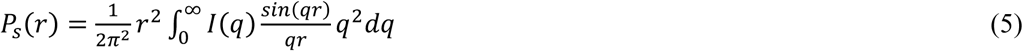

where q is the distance from the origin, P_s_(r) is a histogram of the lengths of interatomic vectors within the scattering object as conventionally calculated from SAXS data from globular proteins. The subscript ‘s’ is introduced here to distinguish it from the cross-section pair-distribution function defined in the next paragraph. The correlation function, a_s_(r) is related to P_s_(r) by

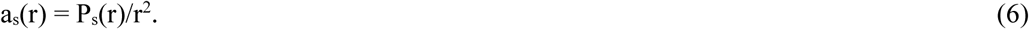

In analysis of scattering from tissues the correlation function has advantages over the pair-distribution function because it suppresses the impact of long range inter-atomic vectors that are due either to the length of the fibril or the spatial correlations between the fibrils of interest and the surrounding polymeric matrix. Although these terms may well provide information about tissue structure within the lesions, they are prone to high uncertainty, very sensitive to the smallest q for which intensities can be accurately measured. In scattering from a sample of randomly oriented fibrils with sufficiently short axial repeat, it is possible to recover the equatorial intensity from the observed intensity through multiplication by a geometric correction factor that, except for very small q, is approximately equal to q (Makowski, 1978). For helical assemblies with longer axial repeat (say ∼ ≥ 30 Å) disorientation may lead to a mixing of equatorial and off-equatorial scattering, precluding accurate measurement of intensities on the equator. Equatorial scattering can be analysed in a manner analogous to spherically averaged data, resulting in a ‘cross-section pair-distribution function’, P_c_(r), that is a histogram of the lengths of interatomic vectors *projected onto a plane perpendicular to the fiber axis* (Roig-Solvas and Makowski, 2017):

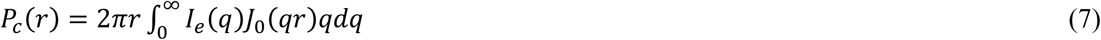

where I_e_(q) is the equatorial scattering, J_o_ is the zeroth-order Bessel function and P_c_(r) is the cross-section pair distribution function. Analogous to the spherical case, a cross-section correlation function, a_c_(r), can be calculated from the cross-section pair distribution function (Glatter, 1980) by

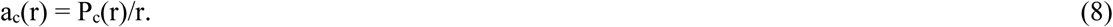

The relationships between these functions are exhibited in **Figure 4** for the case of solid cylinders of length 300 Å or 1000 Å and radius 35 Å. Correlation functions were calculated by an indirect Fourier transform (Glatter, 1980) of spherically averaged intensities (square of Fourier transform) calculated for a solid cylinder of these dimensions and exhibited in **Figure S1**. For calculation of the cross-section correlation function, the equatorial intensity was approximated by multiplying the spherically averaged intensity by a geometric correction factor proportional to q. **Figure 4** demonstrates that these two correlation functions provide direct information on the size and shape of the scattering objects. The maximum vector length for a_s_(r) corresponds to the length of the cylinder; the maximum vector length for a_c_(r) corresponds to its diameter. These functions were computed with data extending to q = 0.3 Å^-1^ so that they correspond to calculations from SAXS data. The limited resolution results in a small over-shoot of zero correlation. As further demonstration of the properties of these functions, the cross-section correlation function was calculated for a model amyloid fibril constructed from 30 layers of Aβ peptides with each layer having the structure of PDB file 5oqv (Gremmer et al., 2017) separated by 4.7 Å from its nearest neighbors. Overall, this structure is ∼ 141 Å long and 56 Å in diameter. As can be seen from **Figure 5** the cross-section correlation function of 5oqv compares well with the correlation function of a 28 Å radius solid cylinder at low r, but deviates from it at higher r due to the non-cylindrical shape of the model amyloid fibril.

**Figure 4.**
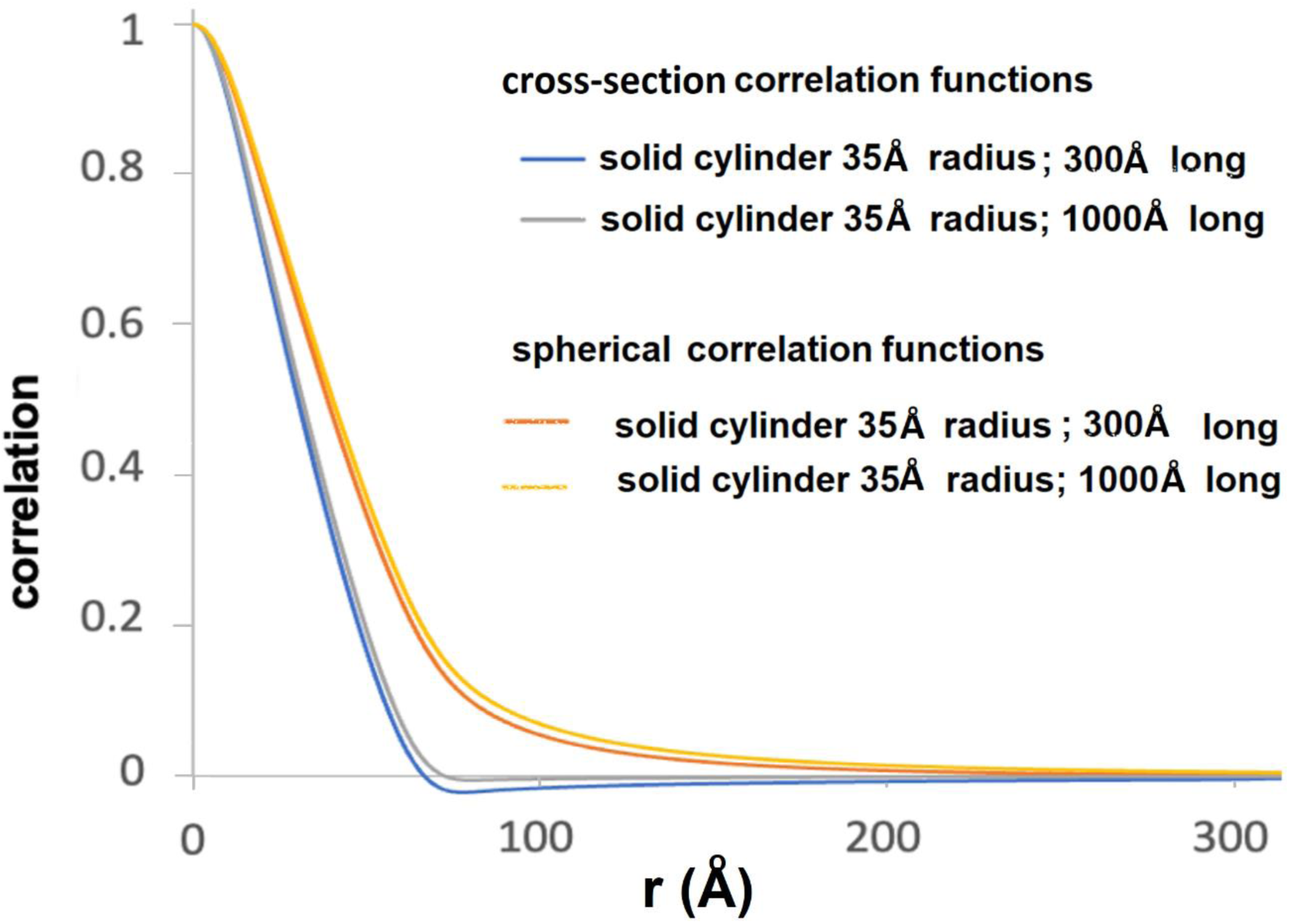
Spherical and cross-sectional correlation functions for solid cylinders 35 Å in radius and 300 or 1000 Å in length calculated from computationally generated intensity distributions. Cross-section correlation functions approach zero at the diameter (70 Å). Spherical correlation functions extend out to a distance equal to their length although for long cylinders its value is quite low for r much greater than the diameter of the cylinder. The slight difference between the cross-section correlation functions calculated for the 300 and 1000 Å in length is due to approximations required to estimate equatorial scattering at very small q.

**Figure 5.**
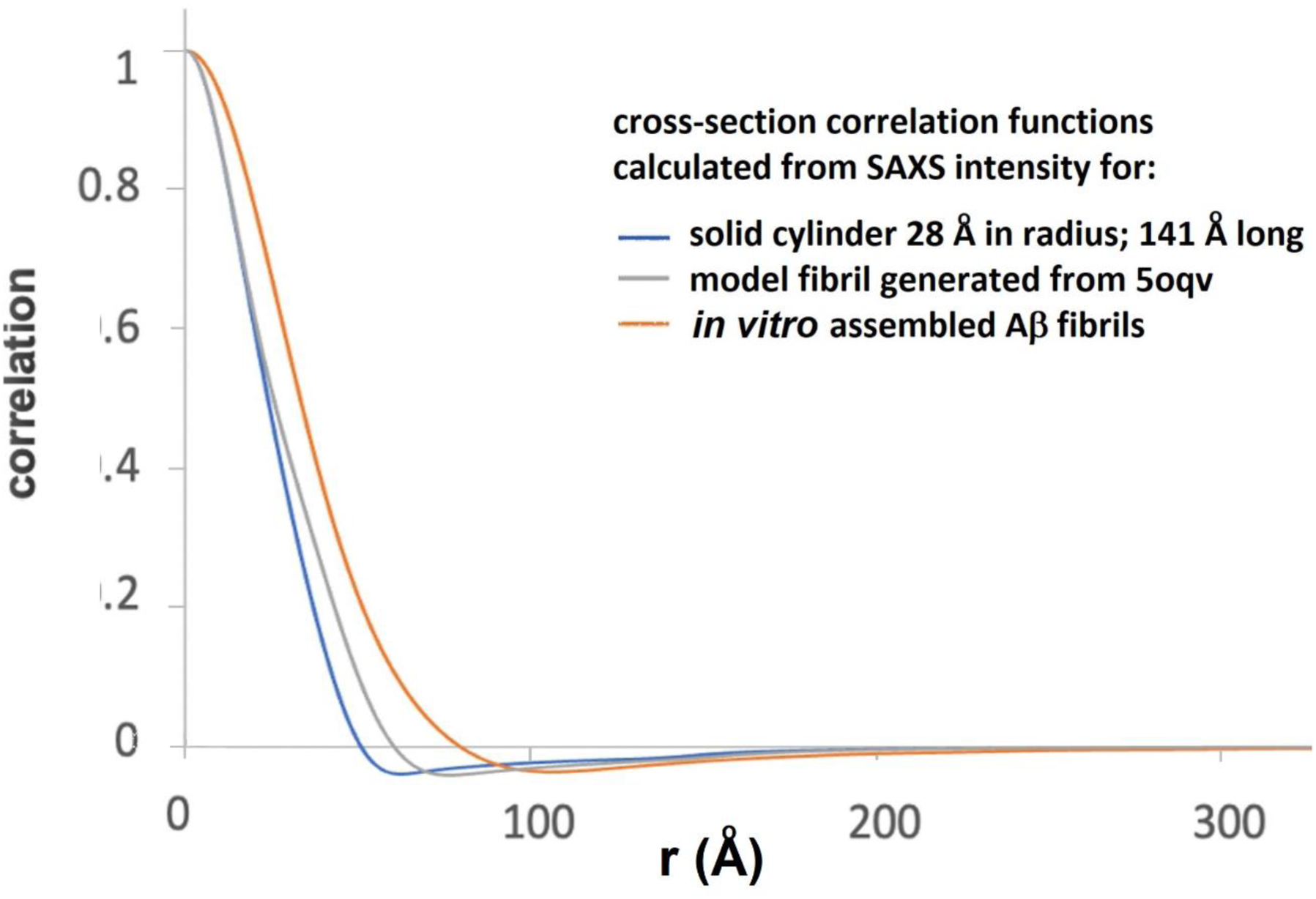
Comparison of cross-section correlation functions from a solid cylinder; a model amyloid fibril generated from pdb file 5oqv and as calculated from SAXS data from *in vitro* assembled Aβ fibrils. At low r, the correlation function of the solid cylinder and the pdb model are indistinguishable since the overall dimensions of the fibril and cylinder are essentially the same. They deviate at larger r due to variation from cylindrical shape in the outer surface of the model fibril. Comparison suggests that the *in vitro* assembled fibrils are significantly larger than the model fibril, consistent with previous modelling of the fibril structure (Roig-Solvas and Makowski, 2017).

### 3.2. Correlation function of Aβ fibrils in aqueous solution

Next, we consider correlation functions calculated from experimental scattering intensities collected from amyloid fibrils in aqueous solution. This is a far simpler environment for scattering than a fixed tissue and it is instructive to consider the scattering in solution as a baseline (Svergun and Koch, 2002; Svergun et. al., 2013) for understanding the observations from scattering when the fibrils are in more complex environments.

We reported SAXS from *in vitro* assembled Aβ peptides (Roig-solvas and Makowski, 2017) and have used those data here to compute cross-section correlation functions. As shown in **Figure 5**, the correlation function calculated from the SAXS data displays properties expected of a fibril with radius greater than that of the model fibril constructed from PDB file 5oqv. We have previously shown that the fibrils in this sample have multiple protofibrils, resulting in a cross-section greater than that of fibril 5oqv and similar to *in vitro* assembled fibrils observed by cryoEM (Meinhardt et al., 2009). The results in **Figure 5** are consistent with those observations.

### 3.3. Correlation functions of solutions and concentrated gels of TMV

SAXS data from increasingly concentrated solutions exhibit features due to interparticle interference and that interference can be characterized through the cross-sectional correlation function. As an example of this, we consider scattering from dilute solutions (Costa et al., 2016) and concentrated gels (Caspar, 1955) of tobacco mosaic virus (TMV). TMV particles are cylindrical rods about 3000 Å long and 180 Å in diameter. Cross-section and spherical correlation functions were calculated from solution scattering data of disoriented dilute solutions provided by Martha Brennich (ESRF) and equatorial scattering from concentrated gels extracted from the thesis of Dr. Donald Caspar (1955) and shown in **Figure S2. Figure 6** displays the results of these calculations.

**Figure 6.**
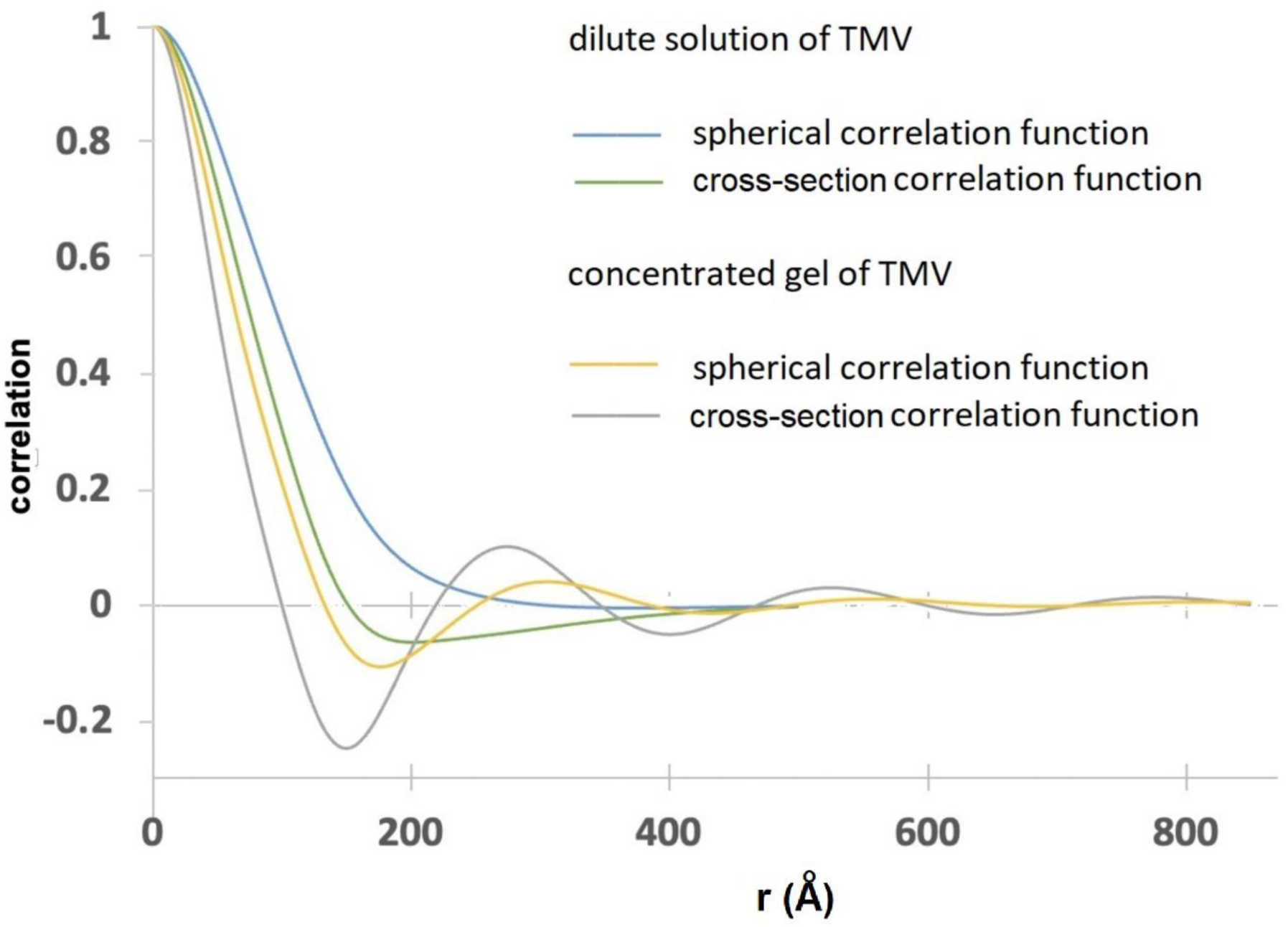
Correlation functions of TMV in dilute solution and concentrated gel.

TMV in concentrated gels is highly oriented making possible accurate measurement of equatorial scattering and observation of dramatic interparticle interference effects (Caspar, 1955). The particles in these gels form a partially ordered, liquid-like two-dimensional packing with nearest-neighbor distances averaging about 275 Å. This is reflected in the peak in the cross-section correlation function in **Figure 6**. This correlation function also exhibits a minimum at about 150 Å at which point the correlation function is considerably below zero. Negative values for correlation functions occur due to regions of electron density below the average electron density of the scattering volume. In these gels, particles have a very low probability of having an interparticle distance of ∼ 150 Å, resulting in an electron density below average for the sample at that radius. This negative correlation is also responsible for the cross-section correlation function crossing zero at r ∼ 100 Å, well below the diameter of the particle, demonstrating that interparticle interference effects can impact estimates of the radius of scattering particles using the cross-section correlation function. The cross-section correlation function calculated from solution scattering approaches zero at about 155 Å, somewhat smaller than expected for the 170-180 Å diameter particles, indicating that even in dilute solution, some interparticle effects are present. The spherical correlation function calculated from dilute solution of TMV approaches zero asymptotically beyond r = 400 Å, reflecting the impact of the particle length on the observed data. While the spherical correlation of the TMV particle should theoretically extend to 3000 Å, intensity measurements were limited to q > 1/400 Å^-1^ and intensity fluctuations due to longer correlations were not observable as indicated in the Methods section.

### 3.4. Correlation function of Aβ fibrils embedded in fixed tissue

**Figure 7** includes the cross-section correlation functions calculated from SAXS data derived from 5 positions in a single dense Aβ plaque embedded in human brain tissue (corresponding to the data shown in **Figure 1** and **Figure S3**). For comparison, the correlation functions of a model fashioned from PDB file 5oqv and as calculated from SAXS scattering from *in vitro*-assembled Aβ fibrils suspended in aqueous solution are included in the Figure. At radii less than the diameter of the fibrils, the cross-section correlation function is dominated by interatomic vectors within the fibril. At intermediate radii, the cross-terms contribute, precluding an accurate estimate of fibril diameter. At radii larger than the diameter, cross-terms between fibril and tissue will dominate. As seen in **Figure 7** the correlation functions from different positions within the dense plaque are nearly indistinguishable at small r, suggesting that the predominant fibrils have the same diameter at each position. They are also nearly identical to the correlation function of the *in vitro* assembled fibrils, but fall at larger radius than calculated from the pdb file 5oqv. This strongly suggests that *in situ*, these fibrils are composed of multiple protofilaments coalesced side-to-side to form larger diameter fibrils. At r >> 30 Å, the correlation function of fibrils within the plaque diverges from the correlation function of the isolated fibrils due to the cross-terms arising from the tissue matrix. The correlation at 200 Å < r < 300 Å varies by 5-10% at different positions in the plaque, suggesting that the mass density varies across the plaque. The correlations within the tissue section may extend to radii larger than that apparent in **Figure 7**, but our ability to estimate their magnitude is limited by the minimum scattering angle for which accurate intensities can be measured. Estimation of correlation functions at radii > ∼ 300 Å are precluded by the x-ray camera geometry used.

**Figure 7.**
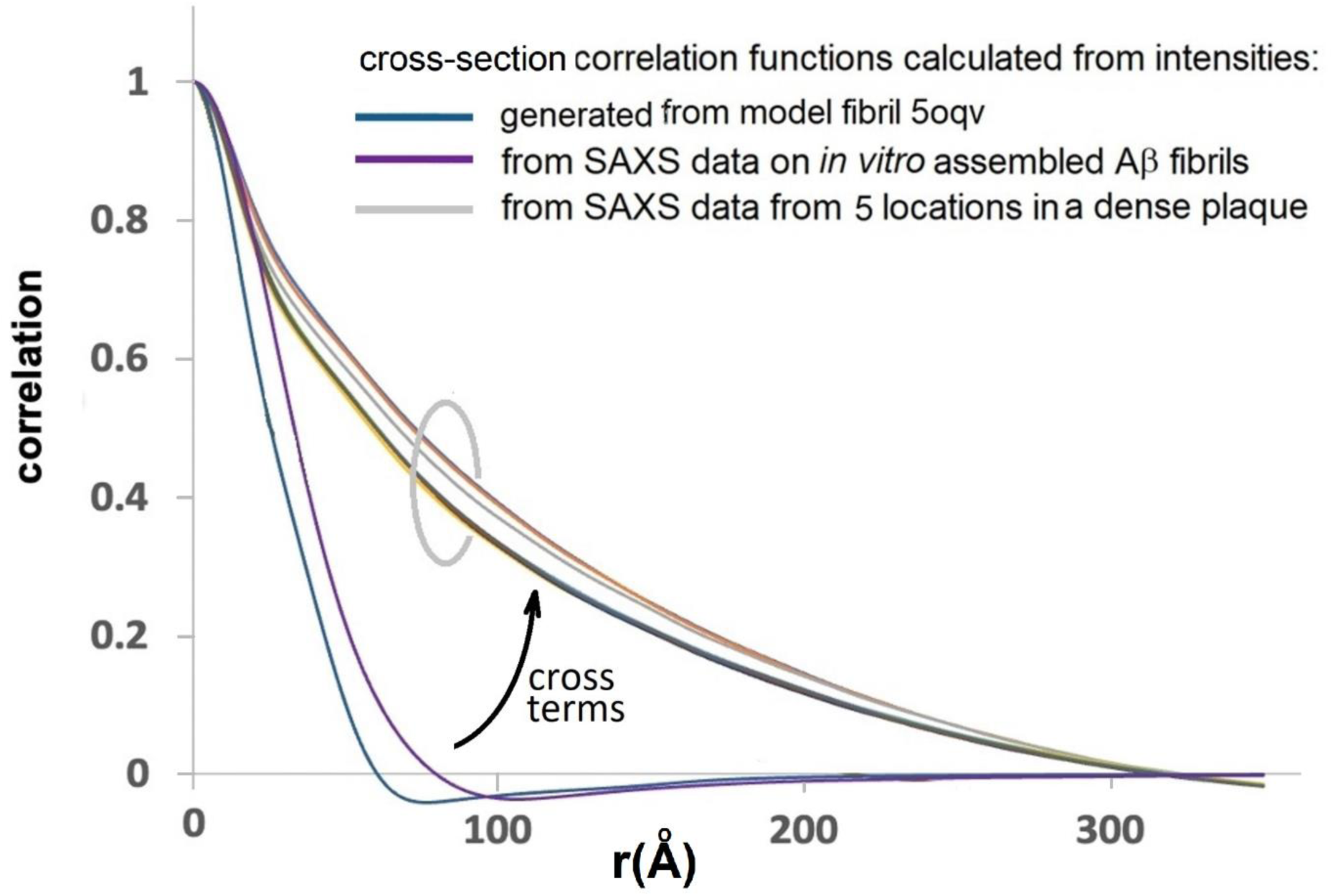
Cross-section correlation calculated for different positions in the dense amyloid plaque detailed in **Figure 1b and 1d**. Compared to correlation functions calculated from SAXS data from *in vitro* assembled fibrils and calculated from a model fibril generated from pdb file 5oqv.

In **Figure 8** the correlation functions of three plaques from the same tissue section are compared as calculated from the data in **Figure S4**. Two of those functions are nearly identical at low radius, suggesting they are dominated by fibrils of identical or similar diameters (by comparison of the red and blue curves at r < 30 Å). The other appears to have a smaller characteristic length (sharper correlation function at low r) and exhibits an oscillation in the correlation with a periodicity of ∼ 50 Å. Analysis of the WAXS data from this lesion suggests it may correspond to a lipidic inclusion or other non-cross-β type structure.

**Figure 8.**
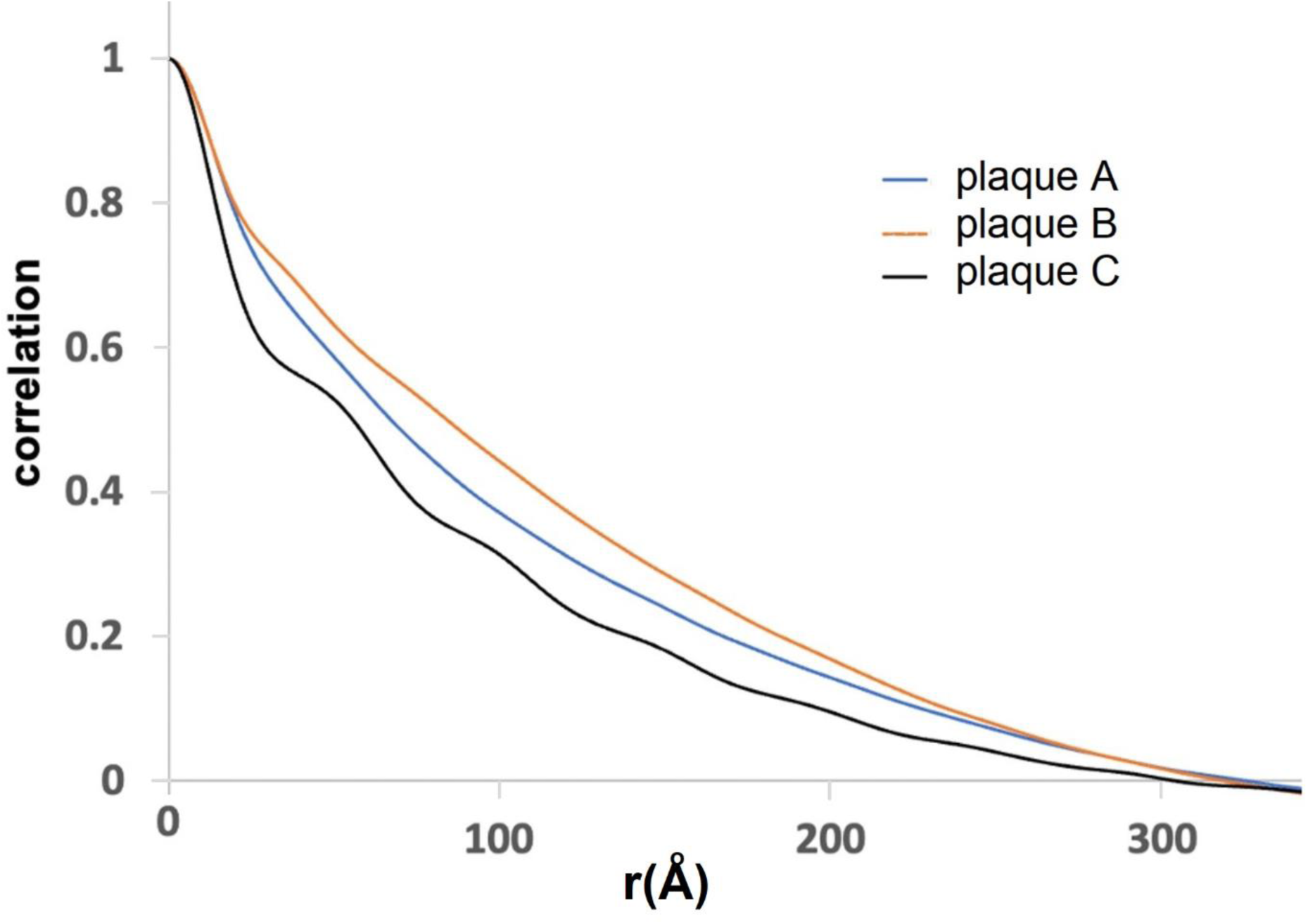
Cross-section correlation functions for three lesions in the tissue section in **Figure 1**. Two plaques (red and blue curves) appear to be very similar at low radius, suggesting similar if not identical fibrillar structure. The third exhibits a low-radius regime suggesting that the predominant scattering particle has a smaller characteristic dimensions.

## 4. Discussion

Scattering from fibrillar structures in solution or embedded in a cross-linked matrix results in data that require special considerations in order to extract useful structural information. In aqueous solution, the scattering from fibrils can be predicted using conventional estimates based on atomic coordinates (Svergun et al., 1995; Park et al., 2009) but their length results in scattering features at very small scattering angles that may be difficult to measure or interpret. When embedded in a solid matrix, the situation is more complex: Spatial correlations between the fibrils and the constituents of the polymer matrix surrounding them will have a significant impact on very small angle scattering. Furthermore, dehydration during sample preparation may increase electron density contrast between the fibrils and their surroundings, thereby enhancing the intensity of small-angle scattering relative to the wide-angle scattering. Depending on the density of the matrix the electron density contrast between fibril and surrounding tissue could be greater in a dehydrated tissue than in dilute aqueous solution, although that is not assured.

In this paper, we described the impact of correlations between fibrils and the other constituents in their immediate vicinity in terms of ‘cross-terms’ in equation (3). An alternative description of these correlations can be made by treating the fibril embedded in tissue as analogous to a protein in aqueous solution. Proteins in aqueous solution are surrounded by a hydration layer in which the water molecules take on a structure distinct from bulk water. To accurately predict SAXS data from the atomic coordinates of these proteins, the correlation between protein atoms and atoms in the hydration layer must be taken into account (Park et al., 2009). This is the strategy we have taken in the formulations described here. However, an alternative formulation is possible in which the hydrated protein is defined as including all protein atoms and all water atoms within the hydration layer. If the hydration layer is chosen to be large enough that its outer layer is indistinguishable from bulk water, then there will be essentially no correlations between the positions of water atoms within and outside the hydration layer. Under those conditions, the ‘cross-terms’ will average to zero. By analogy, an ‘aggregated fibril’ within a tissue matrix might be defined as including both the fibril and the immediately adjacent tissue constituents that take on a distinct organization due to the presence of the fibril (for instance, by binding to or aggregating with the fibril). The advantage of this formulation is that it makes clear that the cross-section correlation function calculated from SAXS of fibrils in tissue describes the properties of the ‘aggregated fibrils’ within the scattering volume. For instance, the ∼ 300 Å radial extent of the cross-section correlation functions plotted in **Figure 7** for an amyloid plaque would indicate that the presence of Aβ fibrils in this plaque impacts the organization of the fixed tissue out to a radius of at least ∼ 150 Å from the fibril axis. The magnitude of correlation at radii greater than the radius of the fibril should provide an estimate of the degree of compaction of material in the immediate vicinity of the fibril (relative to that of bulk tissue). As demonstrated in **Figure 6**, negative values of the correlation are possible if the density of tissue constituents in the immediate vicinity of the fibril is less than bulk tissue.

It is common practice to calculate a pair-distribution function from SAXS data on globular proteins. This is because the domain structure and shape of a globular protein appear well represented in the pair distribution function; and in dilute solutions there is little contribution from inter-protein correlations so there is a well-defined r_max_ beyond which the pair-distribution function is effectively zero. In scattering from fibrils, the conventional SAXS pair-distribution function is far less useful. First, r_max_ is very large (roughly half the length of the fibril), making its estimate highly sensitive to the minimum q for which accurate intensities can be measured. Second, in scattering from fibrils of heterogeneous length, r_max_ becomes ill defined. Third, r_max_ is rigorously the maximum extent of spatial correlations in the sample. For fibrils embedded in a polymeric matrix, the spatial correlations between fibril and surrounding matrix constituents may have very large correlation lengths, corresponding to very large r_max_, the estimation of which will be similarly sensitive to measurement errors in intensities at very small q. To address these challenges, we have utilized the cross-section correlation function (Glatter, 1980). This function is independent of the length of the fibrils and is less sensitive to intensity variation at very small q. It can be calculated for fibrils with axial repeats ≤ ∼ 30 Å for which equatorial intensities can be extracted from SAXS data (where we assume that SAXS data extends to ∼ 30 Å spacing). For fibrils with longer repeat distances, the mixing of equatorial and off-equatorial intensities due to disorientation will limit the maximum scattering angle to which equatorial scattering can be unambiguously measured.

The cross-section correlation function is readily calculated from the cross-section pair-distribution function which constitutes a histogram of interatomic distances *projected perpendicular to the fibril axis*. These cross-section functions are independent of fibril length and have proven useful for fibrils in solution (Roig-Solvas and Makowski, 2017). In scattering from fibrils embedded in fixed tissue, their form is impacted by the presence of other tissue constituents (as shown above). For these samples, the correlation function is preferred to the pair-distribution function since it is less sensitive to errors in intensity at small q, and to the minimum q for which intensities can be measured.

The spatial correlations between fibril and the surrounding matrix will contribute to the cross-section correlation function calculated from the scattering of fibrils and may preclude an unambiguous estimate of the diameter of the fibrils themselves. The maximum extent of spatial correlations within the tissue matrix may be difficult to estimate since correlations that extend greater than several hundred angstroms will give rise to scattering at very small angles that may not be accessible for some x-ray camera geometries. Nevertheless, the cross-section correlation function provides significant insight into the structure of the fibril-matrix mixture within the scattering volume.

It may be possible to estimate the density of the immediate microenvironment of a fibril from its cross-section correlation function. If that proves successful it will provide insight into the density and structural organization of the pathological lesions in which the fibrils are the predominant species. The way that the cross-section correlation function deviates from that expected from a uniform mass distribution may provide insight into the internal structure of a lesion. We have demonstrated that SAXS data collected from small scattering volumes of fixed tissue when dominated by pathological fibrillar aggregates can provide considerable insight into the structural organization of these lesions - Aβ plaques or neurofibrillary tangles. The variation of these features may provide insights into the way these lesions assemble and grow during the progression of neurodegeneration (Murray et al., 2011).

## Acknowledgements

We would like to acknowledge the considerable contributions of Tessa Connors and Dr. Bradley Hyman at the Massachusetts Alzheimer’s Disease Research Center (MADRC). All tissue samples were prepared by Tessa Connors at (MADRC) in collaboration with Dr. Bradly Hyman.

## Supporting information

**Figure S1.**
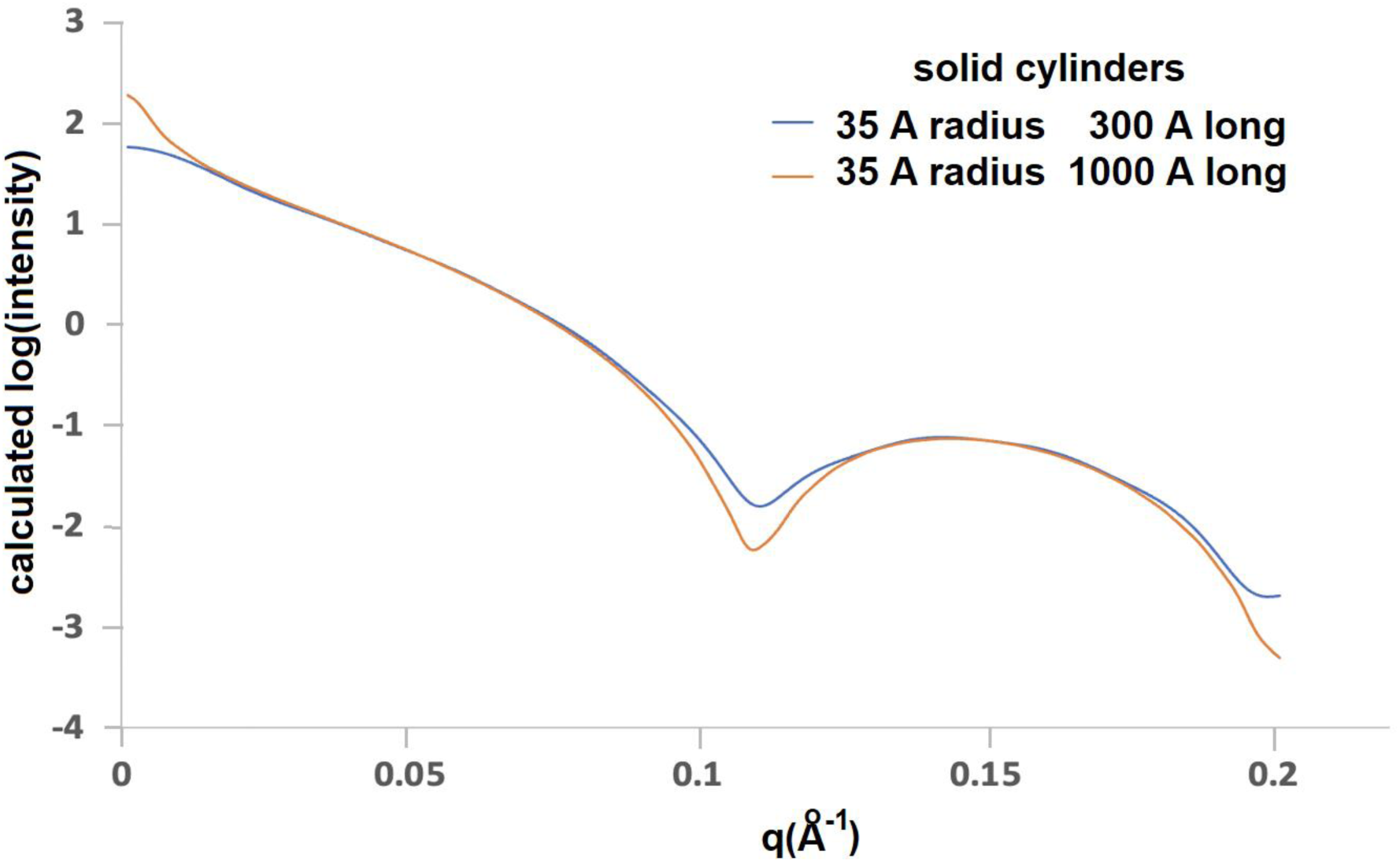
Calculated spherically averaged intensity for solid cylinders 35 Å in radius and 300 and 1000 Å in length. Scattering from the longer cylinder is predicted to exhibit a spike in intensity at very low scattering angles, and sharper minima in intensity.

**Figure S2.**
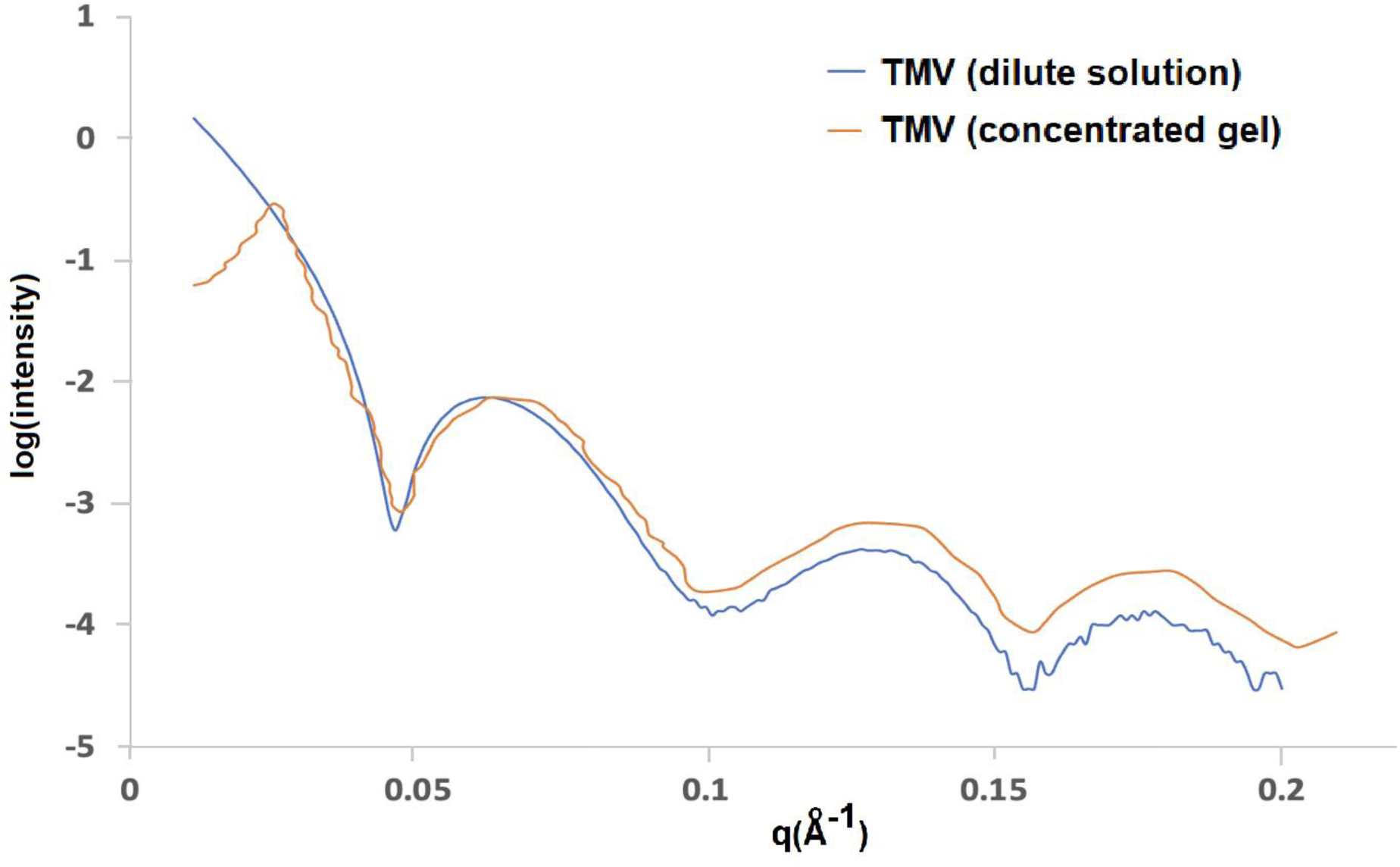
SAXS intensity observed from Tobacco Mosaic Virus in dilute aqueous solution and concentrated gels (after geometric correction). The intensity distributions are very similar except at small angles where interparticle interference effects dramatically alter the scattering from the concentrated gel sample.

**Figure S3.**
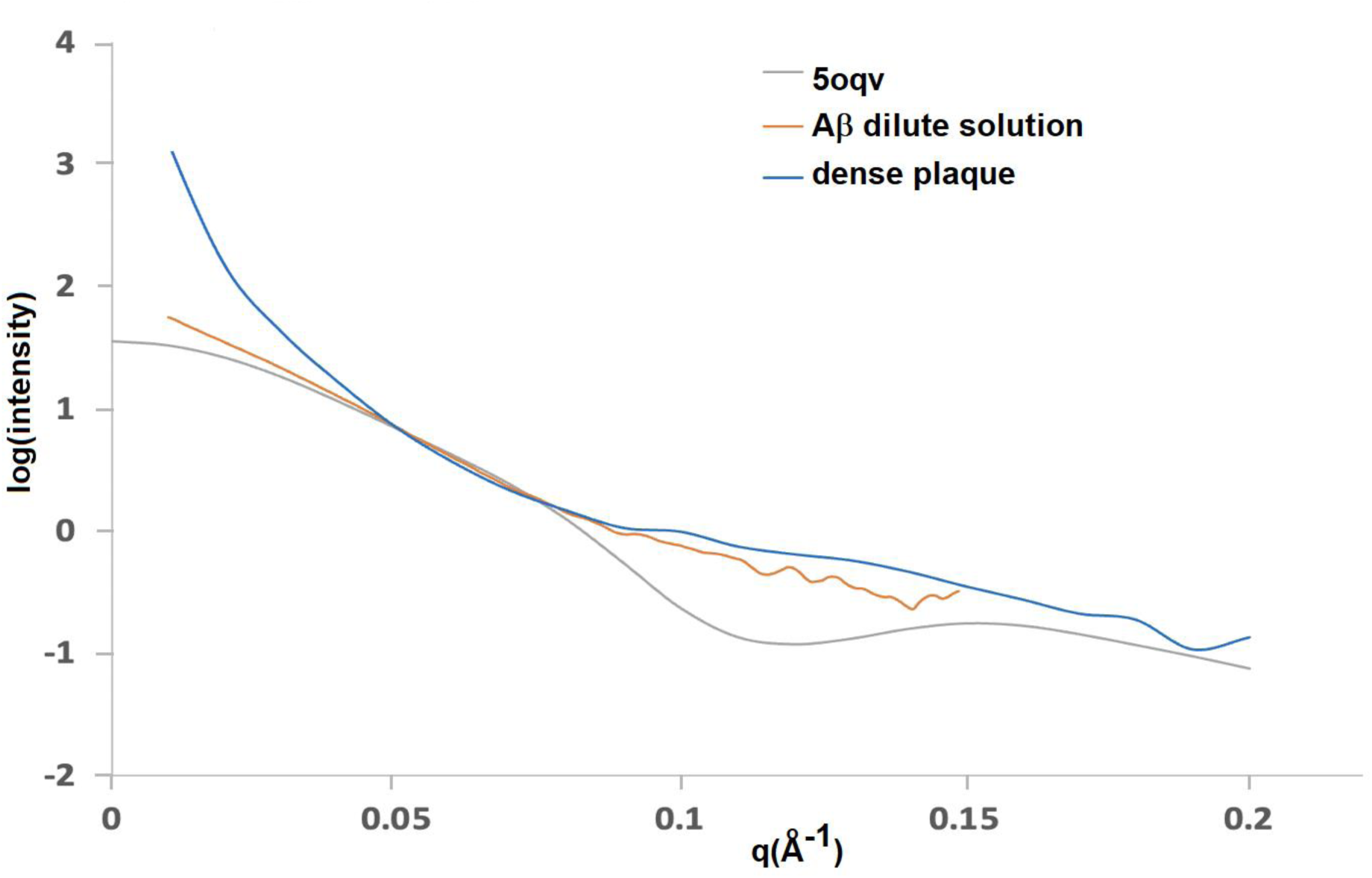
Comparison of SAXS data as estimated from a fibril constructed from pdb file 5oqv; solution scattering from in vitro assembled fibrils; microdiffraction from fibrils embedded in a histological tissue section. Note the small angle spike in scattering from the tissue section.

**Figure S4.**
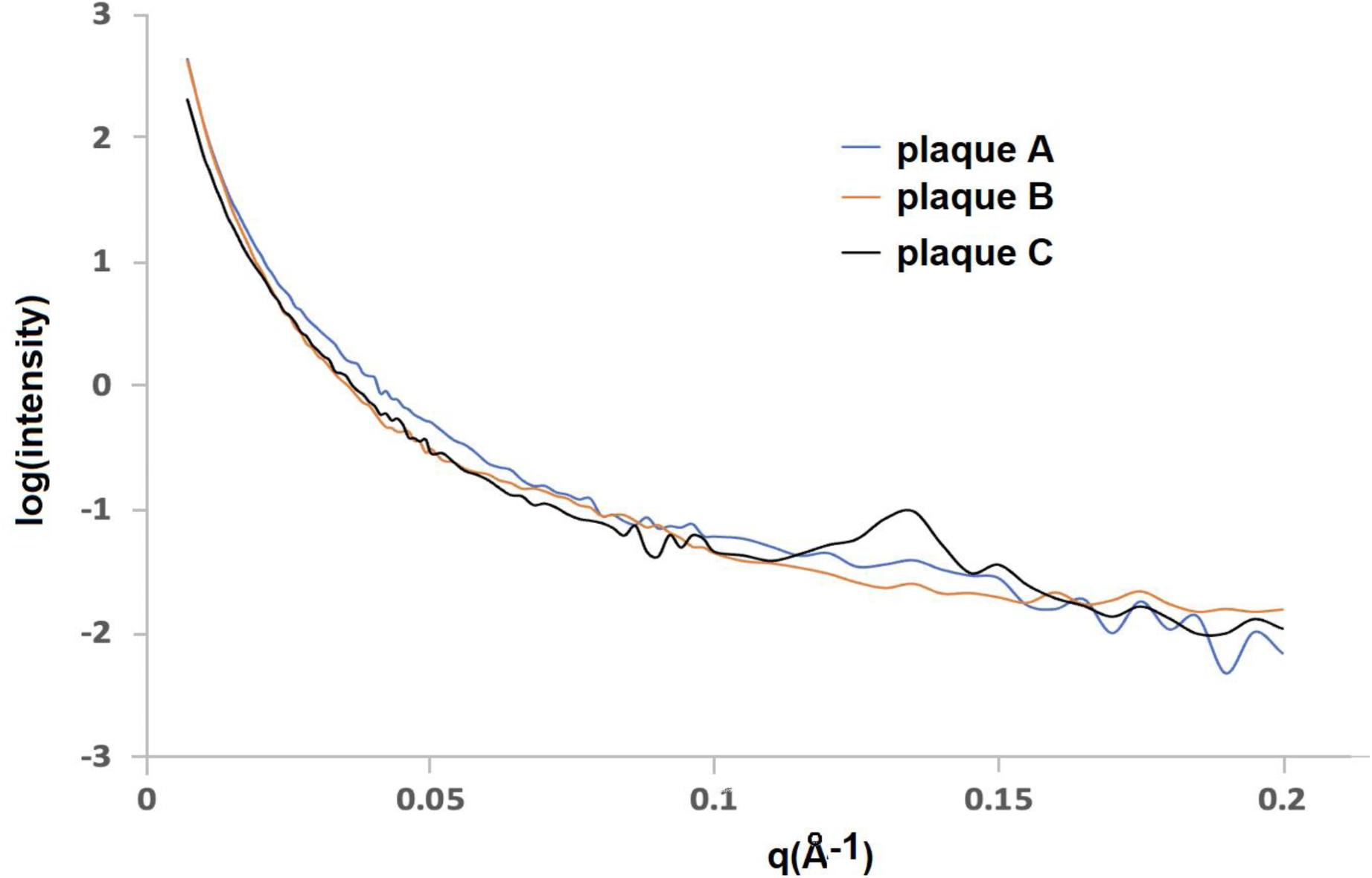
SAXS data from three locations in the sample exhibited in **Figure 1**. Plaque C exhibits a peak at q ∼ 0.14 Å^-1^ corresponding to a periodicity of ∼ 50 Å as seen in the correlation function in **Figure 8**.

